# Deep-learning based retinal fluid segmentation in optical coherence tomography images using a cascade of ENets

**DOI:** 10.1101/2021.09.24.461713

**Authors:** Sophie Loizillon, Cédric Meurée, Camille Breuil, Timothée Faucon, Arnaud Lambert

**Author notes:** These authors also contributed equally to this work (Cédric Meurée, Camille Breuil).

## Abstract

Optical coherence tomography (OCT) is a non-invasive, painless and reproducible examination which allows ophthalmologists to visualize retinal layers. This imaging modality is useful to detect diseases such as diabetic macular edema (DME) or age related macular degeneration (AMD), which are associated with fluid accumulations. In this paper, a cascade of deep convolutional neural networks is proposed using ENets for the segmentation of fluid accumulations in OCT B-Scans. After denoising the B-Scans, a first ENet extracts the region of interest (ROI) between the inner limiting membrane (ILM) and the Bruch’s membrane (BM), whereas the second ENet segments the fluid in the ROI. A random forest classifier was applied on the segmented fluid regions to reject false positive. Our framework was trained on three different datasets with several diseases such as diabetic retinopathy (DR) and AMD. Our method achieves an average Dice Score for fluid segmentation of 0.80, 0.83 and 0.83 on the UMN DME, UMN AMD and Kermany datasets respectively.

## Introduction

Over the last decades, different approaches have been proposed to automatically segment fluid in OCT images. As fluid accumulations are often considered as biomarkers of retinal diseases, their accurate segmentation is crucial for the diagnosis of different pathologies, as well as for the evaluation of the effectiveness of a treatment. Three types of fluid accumulation can occur in the retina: intraretinal fluid (IRF), subretinal fluid (SRF), and pigment epithelial detachment (PED).

Early papers on segmentation in OCT used methods based on thresholds and graph theory. Wilkins et al. [1] proposed an algorithm based on intensity thresholds for fluid segmentation in OCT images. The disadvantage of this approach is that it requires high image quality, which is a well known challenge in the medical imaging field. Rashno et al. [2] used the graph cut method to segment fluid accumulations. These classical image processing algorithms have the inconvenience of a high computational time, which did not allow ophthalmologists to include them in their daily workflow.

These aspects, together with the development of the field of machine learning over the last decades, have led to the publication of papers on fluid segmentation in OCT based on machine learning methods. Random forest classification [3], kernel regression [4] and fuzzy level set [5] methods have been implemented to perform fluid segmentation. These methods involve the training of a classifier by extracting a large number of textural, structural or positional features from OCT images. Machine learning approaches have allowed significant improvements in segmentation results over classical image processing.

In recent years, deep learning methods have been more specifically used in medical image processing. The U-Net architecture presented by Ronnenberger et al. [6] has provided a real breakthrough in biomedical image segmentation. Inspired by the Fully Convolutional Network (FCN), U-Net combines deep semantic and spatial information through encoding and decoding blocks linked by skip connections. This architecture has achieved outstanding results in many medical image segmentation tasks and has been used for OCT segmentation, whether for fluid region segmentation [7], retinal layer segmentation [8], drusen segmentation [9], or intraretinal cystoid fluid segmentation [10]. Lu et al. [7], winners of the Retinal OCT Fluid Challenge (RETOUCH), used a U-Net to segment the three different types of fluid in B-Scans. Venhuizen et al. [10] proposed a cascade of two U-Nets with one extracting the region of interest and the second segmenting the fluid regions. Chen et al. [11] integrated squeeze and excitation (SE) blocks into the classical U-Net structure to only keep the useful feature maps for fluid regions segmentation. Roy et al. [12] implemented a new fully convolutional architecture inspired by U-Net, called ReLayNet, to segment both retinal layers and fluid accumulations in OCT images. Other FCN-based architectures such as Seg-Net and Deeplabv3+ have also been used for fluid segmentation in B-Scans showing good results [13] [14].

Recently, the Efficient Neural Networks (ENet) architecture, which was initially developed for real-time semantic segmentation, has been trained on biomedical images for the segmentation of aneurysms, prostate and skin lesions [15] [18]. ENet contains only half the number of parameters of the U-Net architecture for similar performances. To our knowledge, ENet has not yet been used for fluid segmentation in OCT B-Scans.

In this paper, we present a novel approach to automatically segment and quantify fluid in OCT B-Scans. Our pipeline is composed of a preprocessing step which uses the BM3D algorithm to reduce the speckle noise. Then, a cascade of deep convolutional neural networks using ENets is trained to extract the region of interest (ROI) and to segment the fluid pixels in the ROI mask. To complete our segmentation pipeline, we refine the results with a post-processing step by using a random forest classifier.

This article is organized as follows: the research materials and methods are described in the second section, including the details of our pipeline implementation and its application to the automatic segmentation of retinal fluids in OCT images. The third section is dedicated to the segmentation and quantification results. Finally, the fourth and fifth sections present the discussions and conclusions, respectively.

## Materials and methods

### Datasets

For the present study, three public OCT datasets were employed to train and test our method. Two of them were created by the University of Minnesota (UMN) [2] [20]. The OCT volumes were acquired with a Heidelberg Spectralis imaging system for 29 subjects with DME and 24 with AMD. Each B-Scan averages 12 to 19 frames with a resolution of (5.88 × 3.87) µm/pixel. The accumulations of fluid were manually segmented by two UMN ophthalmologists.

The Kermany dataset consists of 530 OCT volumes divided into three categories: DME, DRUSEN and NORMAL [21]. The images were also acquired with a Heidelberg Spectralis imaging system and manual fluid segmentation was performed by three trained graders and approved by two ophthalmologists. For each volume, the number of B-Scans varied from 1 to 13, with most volumes containing only one or two.

### Segmentation Pipeline

Our pipeline for retinal fluid segmentation and quantification in OCT images consists of five steps, as shown in Fig 1: 1) preprocessing of the OCT B-Scans by means of the BM3D algorithm; 2) segmentation of the ROI located between the ILM and the BM; 3) segmentation of the pixels associated with fluid accumulations using a convolution neural network; 4) post-processing of the binary fluid segmentation mask using a random forest classifier; 5) 2D fluid surface quantification.

**Fig 1.**
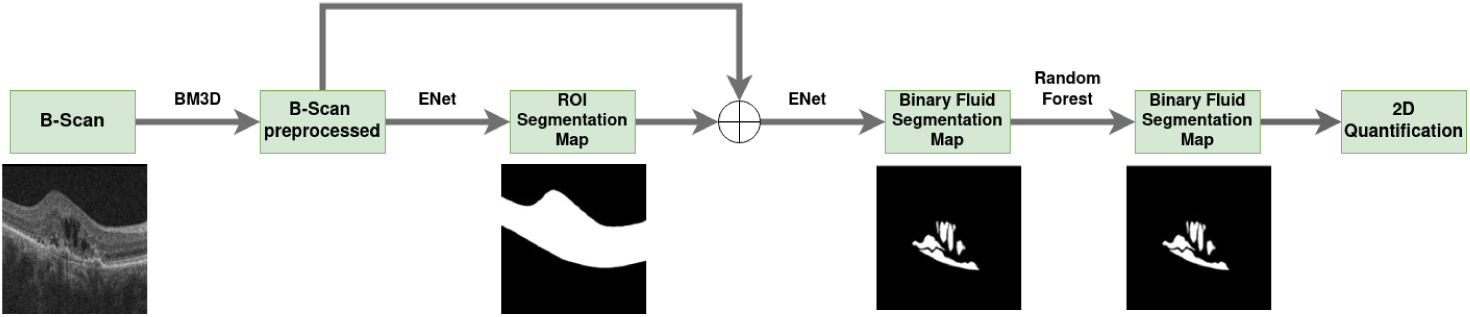
Proposed method for fluid segmentation and quantification in OCT B-Scans.

#### Preprocessing

Speckle noise is the main factor which degrades the quality of OCT images. This granular and multiplicative noise impacts the performance of automatic analysis in OCT images. Based on the literature, we investigated three algorithms in order to reduce the speckle noise in B-Scans. We implemented the Block Matching and 3D Filtering (BM3D) algorithm proposed by Dabov et al [17], which is one of the most powerful image denoising methods with its collaborative filtering. We also tested the Non Local Means (NLM) algorithm presented by Buades et al. [19] which exploits the presence of similar features within an image. For the NLM and BM3D, we used a seach window of 21 pixels and a patch size of 7. Finally, we implemented the median filter (as 1D filter of 5 pixels).

#### ROI Segmentation

Based on Venhuizen et al. [10] method, we decided after having denoised the B-Scans to perform a segmentation between the ILM and BM layers, where fluid accumulation can be located. We investigated four architectures encountered in the literature: the U-Net, SEU-Net, Seg-Net and ENet in order to identify the most efficient one for the ROI segmentation in OCT B-Scans. Each architecture takes as input a B-Scan preprocessed by the BM3D algorithm and generates as output a binary segmentation mask of the ROI. This step has the advantage of removing all structures below the BM, including the choroid, which can degrade the automatic analysis of OCT images.

#### Fluid Segmentation

The second model of this pipeline generates a binary fluid segmentation mask. It takes as input two images: the B-Scan preprocessed by the BM3D algorithm and the output of the first model, namely the ROI mask. One of the major advantages of this approach is that the model will be able to focus only on pixels that are likely to contain fluid. For this segmentation task, we also tested the same four architectures to identify the most relevant one. We compared our performances with several approaches encountered in the literature: Rashno et al. [2] and Ganjee et al. [16] methods for the UMN DME dataset, Rashno et al. [2] and Chen et al. [11] for the UMN AMD, and Ganjee et al. [16] and Lu et al. [7] for Kermany.

#### Post-processing

After training our cascade of deep convolutional neural networks to segment fluid in OCT images, we added a post-processing step to our segmentation pipeline. This step improves the performance of our method by rejecting false positive detected pixels. Three machine learning classifiers were identified in the literature: the random forest, the support vector machine and the K-nearest neighbour methods. Each one was trained by extracting the following properties from potential fluid regions: perimeter, area, average intensity, variance of the intensity, orientation and eccentricity.

#### Quantification

Different machines are capable of acquiring OCT images. These can have different resolutions. Thus, a quantification step is needed to compare segmentation results. Also, most ophthalmologists use quantitative OCT biomarkers, such as the amount of fluid accumulations to inform treatment decisions in individualized therapies. This is why we decided to implement a 2D quantification step in our pipeline. To quantify the surface of fluid in B-Scans, we first evaluate the number of detected fluid pixels thanks to the segmentation map and then calculate the product between the number of detected pixels and the pixel resolution.

### Implementation

Our proposed method is a cascade of deep convolutional neural networks composed of two ENets. It was originally developed for real-time semantic segmentation and the architecture contains half the number of parameters of a U-Net for similar performances. The ENet architecture is divided into five stages, three of which are dedicated to the encoder and two to the decoder. Each stage is composed of convolution blocks with short skipped connections known as bottleneck blocks. They consist of three convolutional layers. A first 1 × 1 projection is used to reduce the dimensionality. Then, a regular, dilated, or full convolution is performed. Finally, a 1 × 1 projection expands the dimensionality of the image. Paszke et al. [15] decided to use an asymmetric architecture because they believed that the encoder should be able to operate on lower resolution than classification architectures.

The two ENets were trained using the Focal Tversky Loss (FTL) function defined in Eq 1. This loss function was used to address the issue of data imbalance for fluid segmentation in OCT images and thus to achieve a better trade off between precision and recall.

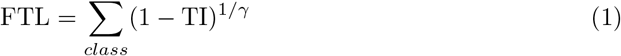

The Tversky Index (TI) is a generalization of the dice score allowing more flexibility in the balance between false positives and false negatives by means of the two scalar hyperparameters *α* and *β* as shown in Eq 2.

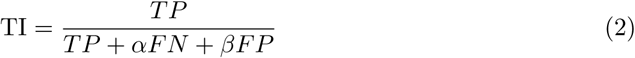

where TP represents the true positive, FN the false negative and FP the false positive pixels.

In Eq 1, *γ* ranges from 1 to 3. We trained both ENets with *γ* > 1 so that the loss function focuses more on less accurate pixel predictions. We performed an optimization of hyperparameters including the *α, β* and *γ* coefficients by Random Search and obtained the best results with *α* = 0.6, *γ* = 0.4 and 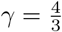 Indeed, using a higher value of *α* improves the model performances by minimizing false negative predictions.

Thanks to the Random Search optimization, we set the learning rate of the Adam optimizer at 10^−4^, the batch size at 32 and the spatial dropout rate of each bottleneck at 0.12. The training process was stopped if the FTL didn’t decrease during 10 epochs.

This segmentation pipeline was implemented using Python and the deep learning API Keras. We used the scikit-learn library for median filtering, the NLM algorithm and the post-processing machine learning classifiers. The BM3D library was also utilized to preprocess the B-Scans.

### Performance Analysis

#### Preprocessing Metrics

To quantify the quality of the preprocessed images, we evaluated several metrics which can be divided into 2 groups: the intensity-based metrics which operate only on the intensity of the distortions and the feature-based metrics, which measure quality based on information or structures from the image. We utilized the Mean Square Error (MSE) and the Peak Signal-to-Noise Ratio (PSNR) as intensity-based metrics. For the feature-based metrics, we have chosen to focus on: the Structural Similarity Index (SSIM), the Multi-scale Structural Similarity Index (MS-SSIM), the Visual Information Fidelity (VIF) and the Universal Image Quality Index (UQI).

#### Segmentation Metrics

To evaluate the performance of fluid segmentation we used the Dice Similarity Coefficient (DSC), which measures the similarity between the predicted segmentation and the ground truth. The formula of the DSC is given in Eq 3.

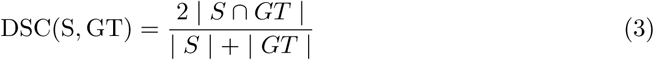

with S the segmentation result and GT the ground truth.

Three other metrics were evaluated as secondary metrics to validate our segmentation models: the Intersection over Union (IoU), Precision and Recall.

Intersection over Union (IoU) evaluates the area of overlap between the predicted segmentation and the ground truth divided by the area of the union between both segmentation.

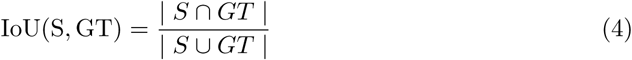

Precision describes the purity of our positive detections with respect to the ground truth.

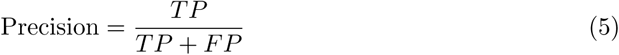

Recall describes the completeness of our positive predictions in comparison to the ground truth.

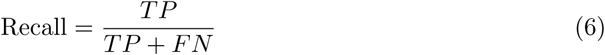

We have also displayed a Receiver Operating Characteristic (ROC) curves for the detection of B-Scans containing fluids between our model predictions and the ground truths. It is a plot of the false positive rate (FPR) (x-axis) versus the true positive rate (y-axis) for a number of different candidate threshold values between 0 and 1. The true positive rate is also known as recall. The false positive rate is defined in Eq 7.

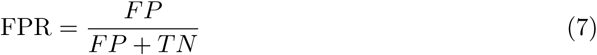

#### Quantification Metrics

To evaluate the 2D quantification performances, we used 2 metrics: Pearson’s correlation coefficient (*ρ*) which reflects a linear relationship between two variables and the coefficient of determination (R^2^) which explains to which extent the variability of a factor can be caused by its relationship with another related factor.

## Results

### Preprocessing

We preprocessed all the B-Scans from the UMN and Kermany datasets with three algorithms: BM3D, NLM and median filtering. The preprocessing results have been summarized in Tab 1.

**Table 1.** Evaluation of three preprocessing algorithms to denoise OCT B-Scans.

|  | MSE | PSNR | SSIM | MS-SSIM | UQI | VIF | Computation time (s) |
| --- | --- | --- | --- | --- | --- | --- | --- |
| NLM | 93 | **32** | 0.68 | 0.91 | **0.85** | 0.24 | 1.9 |
| BM3D | 83 | 34 | 0.67 | 0.90 | 0.84 | 0.25 | 17.23 |
| Median Filtering | **32** | 31 | **0.76** | **0.94** | **0.85** | **0.33** | **0.002** |

The results of the feature-based metrics clearly show that the median filter allows a better preservation of the image features associated with a short execution time, with a SSIM and a VIF 8% higher than the other methods. However, the median filter does not remove as much noise as the BM3D and NLM algorithms. Because of its good ability to remove speckle noise according to its PSNR score, while preserving edges, we decided to preprocess our B-Scans with the BM3D algorithm.

### ROI Segmentation

After having preprocessed the B-Scans with the BM3D algorithm to reduce the speckle noise, we trained four different network architectures to determine the best one for segmenting the ROI. We evaluated the models performances with several metrics as reported in Tab 2.

**Table 2.** Evaluation of four architectures for ROI segmentation in OCT.

|  | DSC | IoU | Recall | Precision | Number of parameters |
| --- | --- | --- | --- | --- | --- |
| U-Net | 0.93 | 0.90 | 0.95 | 0.93 | 7 759 521 |
| SEU-Net | 0.94 | 0.88 | 0.97 | 0.93 | 8 634 913 |
| Seg-Net | 0.93 | 0.89 | 0.95 | 0.92 | 2 932 929 |
| ENet | **0.96** | **0.92** | **0.97** | **0.95** | **371 121** |

These results allowed us to identify the most interesting architecture for ROI segmentation in B-Scans. Indeed, if the difference in DSC remains small between the different architectures, the ENet contains only 371 121 trainable parameters. This makes it possible to significantly speed up the training phase as well as its prediction time. Automatic segmentation of a B-Scan using a ENet architecture takes 0.2 seconds compared to 1.3 seconds for a U-Net.

### Fluid Segmentation

Once we finished the preprocessing step of the OCT B-Scans with the denoising algorithm BM3D and the ROI segmentation, we were able to perform fluid segmentation. As we previously did for the ROI, we determined the best possible architecture for segmenting fluid accumulations in OCT images. Therefore, we trained four different architectures using the preprocessed and ROI segmented B-Scans from the three datasets. Results are detailed in Tab 3.

**Table 3.** Evaluation of four architectures for fluid segmentation in OCT.

|  | DSC | IoU | Recall | Precision |
| --- | --- | --- | --- | --- |
| U-Net | 0.76 | 0.66 | 0.77 | 0.78 |
| SEU-Net | 0.78 | 0.69 | 0.79 | 0.80 |
| Seg-Net | 0.79 | 0.69 | 0.80 | 0.82 |
| ENet | **0.81** | **0.70** | **0.86** | **0.89** |

Thanks to this comparative analysis, we were able to identify the most interesting network architecture for fluid segmentation in OCT B-Scans. Once again, the ENet architecture gave us the best DSC with 81%. Fig 2 shows the segmentation results obtained with the four different network architectures on a B-Scan from the UMN DME dataset.

**Fig 2.**
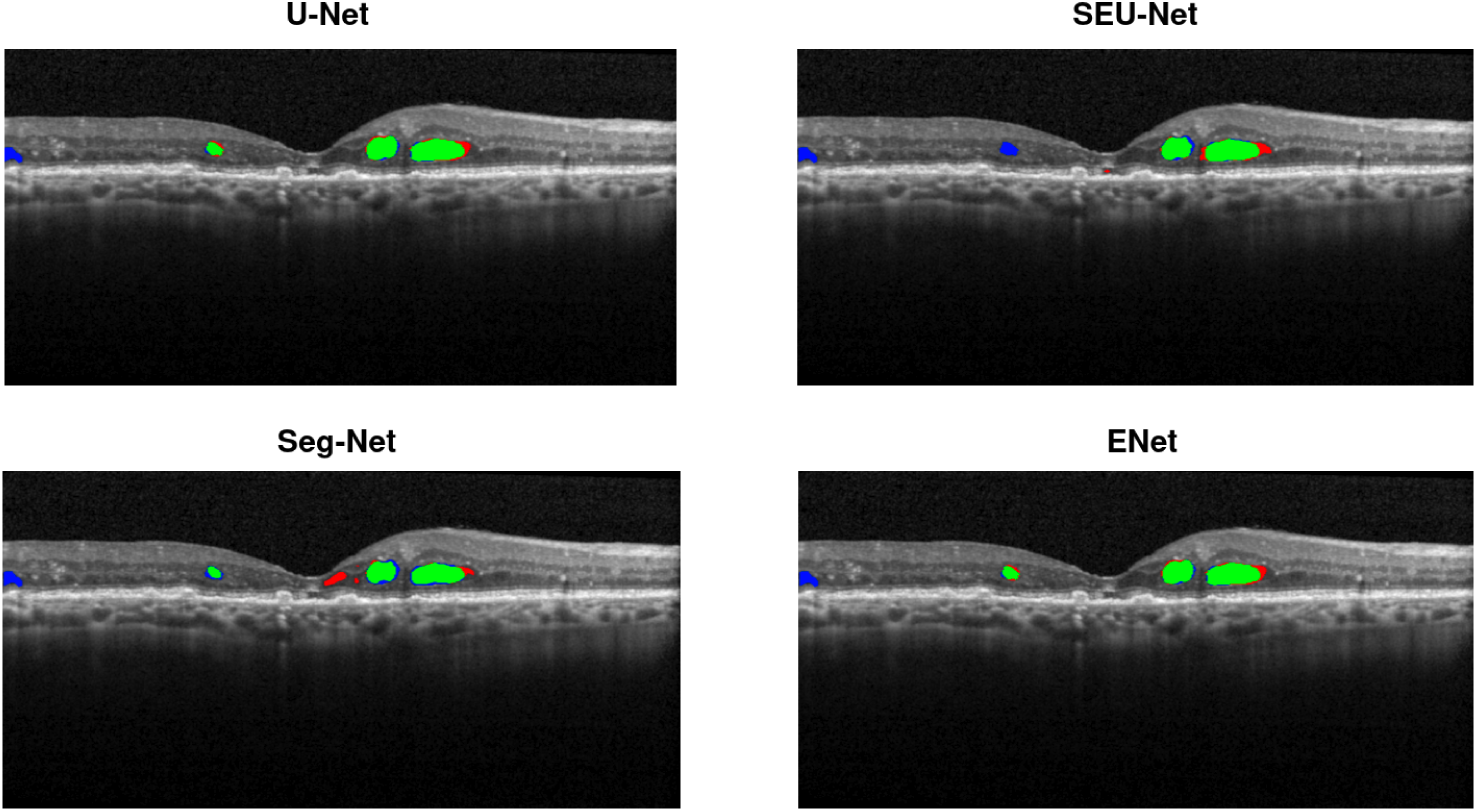
Fluid regions segmentation results using different network architectures. Segmentation maps with green pixels correspond to true positives, red to false positives and blue to false negatives

### Post-processing

We completed our segmentation pipeline with a post-processing step to reject false positives. We tested three machine learning classifiers: the random forest, the support vector machine and the k-nearest neighbors as shown in Tab 4

**Table 4.** Evaluation of four architectures for fluid segmentation in OCT.

| Architecture | Post-processing | DSC | IoU | Recall | Precision |
| --- | --- | --- | --- | --- | --- |
| ENet | – | 0.81 | 0.70 | 0.86 | 0.89 |
|  | RF | **0.82** | **0.70** | **0.87** | **0.90** |
|  | SVM | 0.80 | 0.69 | 0.85 | 0.88 |
|  | k-NN | 0.81 | 0.70 | 0.87 | 0.90 |

Thus, the addition of a post-processing step based on the random forest machine learning classifier allowed us to gain 1% of Dice Score. This step rejects several false positives and therefore increases the recall score as shown in Fig 3.

**Fig 3.**
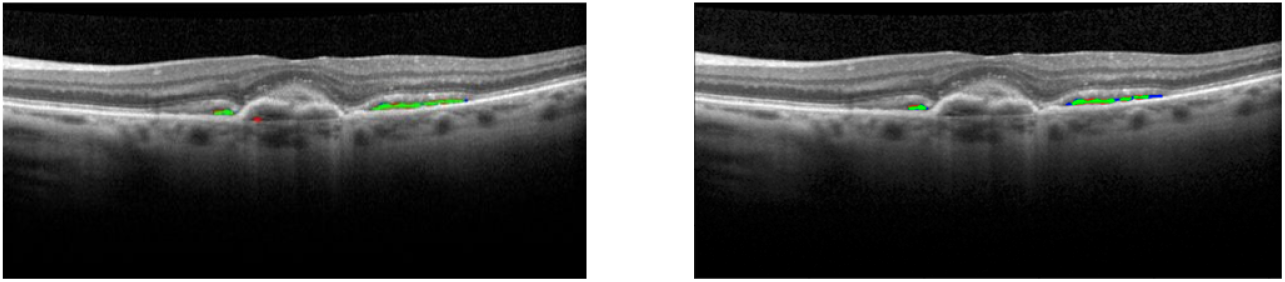
Segmentation map without and with a post-processing step. On the left the first segmentation map without any post-processing step and on the right the post-processed segmentation map with a random forest classifier.

### Full Segmentation Pipeline

Once the models architecture was selected and the pipeline implemented, we investigated its capability to generalize fluid segmentation for several diseases. The DSC boxplot and the ROC curves of fluid detection are displayed in Fig 4. For each dataset, the AUC was above 0.97, indicating an excellent detection accuracy regardless of the type of pathology.

**Fig 4.**
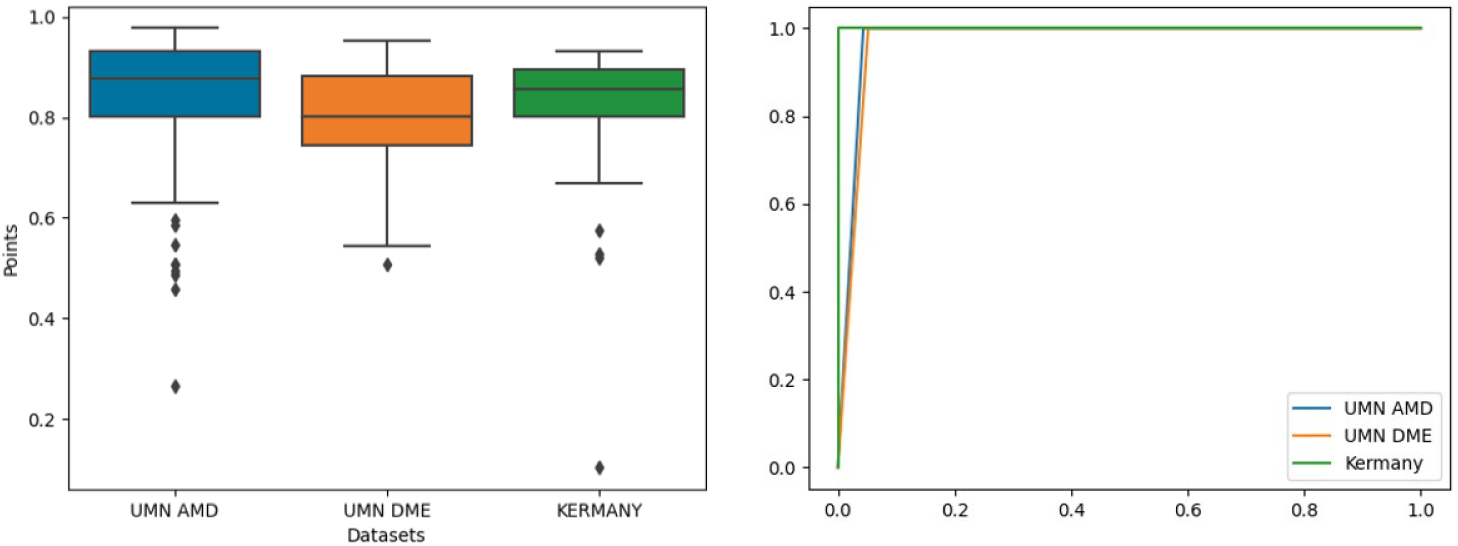
DSC boxplot for fluid segmentation and ROC curves for fluid detection of three different datasets.

Finally, we compared our pipeline performances with approaches encountered in the literature as shown in Tab 5. We observed an improvement of over 10% and 5% on Rashno et al. [2] and Ganjee et al. [16] methods respectively for the UMN DME dataset. A smaller improvement was obtained for the Kermany dataset with a gain of 4% and 1% compared with Ganjee et al. [16] and Lu et al. [7]. However, our pipeline did not reach the state of art performance of Chen et al for the fluids segmentation on the UMN AMD dataset.

**Table 5.** Comparison of Dice Similarity Coefficient results for the segmentation of fluid in OCT.

| Dataset | Paper | Methods | DSC |
| --- | --- | --- | --- |
| UMN DME | Rashno et al. | Unsupervised | 0.69 |
|  | Ganjee et al. | Unsupervised | 0.75 |
|  | **Our Method** | **Supervised** | **0.80** |
| UMN AMD | Rashno et al. | Unsupervised | 0.82 |
|  | **Chen et al.** | **Supervised** | **0.94** |
|  | Our Method | Supervised | 0.83 |
| KERMANY | Ganjee et al. | Unsupervised | 0.79 |
|  | Lu et al. | Supervised | 0.82 |
|  | **Our Method** | **Supervised** | **0.83** |

### Quantification

We evaluated the performance of our fluid quantification step using two metrics: the Pearson correlation coefficient *ρ* and the coefficient of determination *R*^*2*^. We could not estimate the fluid surface for the Kermany dataset, because the resolution of the B-Scans was unavailable. For the UMN datasets, comparison with assessment by expert graders trained in retinal analysis showed a good correlation with automated quantification of fluid in B-Scans as reported in Tab 6.

**Table 6.**
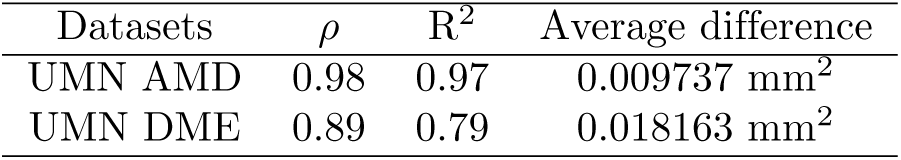
Evaluation of the quantification of fluid in OCT images.

| Datasets | $\rho$ | $R^2$ | Average difference |
| --- | --- | --- | --- |
| UMN AMD | 0.98 | 0.97 | 0.009737 mm<sup>2</sup> |
| UMN DME | 0.89 | 0.79 | 0.018163 mm<sup>2</sup> |

## Discussion

We present a fluid segmentation method on OCT B-Scans based on a cascade of deep convolutional neural networks using ENets. To evaluate the performance of the proposed method, we have compared it to the fluid segmentation methods encountered in the literature. As shown in Tab 3, the proposed method achieves good results on OCT B-Scans related to DME and AMD affected patients. On the UMN DME dataset, we outperform the unsupervised approaches of Rashno et al. [2] and Ganjee et al [16] by up to 10% for the dice score. Both papers proposed unsupervised approaches and therefore tested their method on the whole dataset. They detailed their results for each volume by taking the mean Dice Score of each B-Scan composing the volume. Thus, we were able to compare our performance on our test set. Our method slightly improved the results obtained by Ganjee et al. [16] with their unsupervised approach on the Kermany dataset and matched the performance of Lu et al. [7]. However, this comparison must integrate the fact that the dataset was unknown to Lu et al. as their model was trained on another dataset provided by the Medical Image Computing and Computer Assisted Intervention conference (MICCAI). We also evaluated our network on AMD OCTs. We compared our performance on the 5 OCT volumes of our test set with the one of Rashno et al [2] and found a very slight improvement in results of 1%. We had more difficulties to compare our results to those of Chen et al. [11], who report a mean Dice Score of 94%, because it was not made clear which OCT volumes were considered in the training and testing phases of their work. To avoid these issues, we have detailed our results for each volume to facilitate future comparisons in Tab 7. A possible explanation for this discrepancy may be that Chen et al. limit themselves to fluid segmentation in AMD, whereas our model seeks to generalize fluid segmentation to different retinal pathologies.

**Table 7.** Mean Dice Score per volume for UMN AMD and DME test set.

|  | UMN DME |  |  |  |  |  | UMN AMD |  |  |  |  |
| --- | --- | --- | --- | --- | --- | --- | --- | --- | --- | --- | --- |
| Subject | 1 | 2 | 3 | 4 | 5 | 6 | 1 | 2 | 3 | 4 | 5 |
| Mean Dice | 0.84 | 0.82 | 0.81 | 0.76 | 0.78 | 0.82 | 0.86 | 0.89 | 0.85 | 0.82 | 0.68 |

In order to assess the generalization and potential clinical application of the proposed framework, it would be necessary to conduct in the future an additional experiment on a clinical routine dataset.

## Conclusion

In this paper, we have described a novel approach to automatically segment fluid in OCT B-Scans. The proposed pipeline starts with a preprocessing step to reduce the speckle noise thanks to the BM3D algorithm. Then, it consists of a cascade of ENets where the first one extracts the region of interest between the ILM and BM and the second one segments fluid accumulations. We complete our network with a post-processing step by training a random forest classifier to remove false positive pixel detections and thus to improve our segmentation performances. The proposed method showed good performances with a DSC over 80% on three different datasets associated with different types of diseases. In the future, we plan to test our method on a “real life” dataset to assess the generalization and clinical benefit of the proposed framework.

## References

1. Wilkins G, Houghton O, Oldenburg A Automated et al. Segmentation of Intraretinal Cystoid Fluid in Optical Coherence Tomography. IEEE Journals & Magazine. 2012; doi:10.1109/TBME.2012.2184759

2. Rashno A, Nazari B, Koozekanani DD, Drayna PM, Sadri S, Rabbani H, et al. Fully-automated segmentation of fluid regions in exudative age-related macular degeneration subjects: Kernel graph cut in neutrosophic domain. PLOS ONE. 2017;12(10):e0186949. doi:10.1371/journal.pone.0186949.

3. Lang A, Carass A, Swingle EK, Al-Louzi O, Bhargava P, Saidha S, et al. Automatic segmentation of microcystic macular edema in OCT. Biomedical Optics Express. 2015;6(1):155–169. doi:10.1364/BOE.6.000155.

4. Chiu SJ, Allingham MJ, Mettu PS, Cousins SW, Izatt JA, Farsiu S. Kernel regression based segmentation of optical coherence tomography images with diabetic macular edema. Biomedical Optics Express. 2015;6(4):1172. doi:10.1364/BOE.6.001172.

5. Wang J, Zhang M, Pechauer AD, Liu L, Hwang TS, Wilson DJ, et al. Automated volumetric segmentation of retinal fluid on optical coherence tomography. Biomedical Optics Express. 2016;7(4):1577–1589. doi:10.1364/BOE.7.001577.

6. Ronneberger O, Fischer P, Brox T. U-Net: Convolutional Networks for Biomedical Image Segmentation. International Conference on Medical Image Computing and Computer-Assisted Intervention. 2015;9351:234–241

7. Lu D, Heisler M, Lee S, Ding GW, Navajas E, Sarunic MV, et al. Deep-learning based multiclass retinal fluid segmentation and detection in optical coherence tomography images using a fully convolutional neural network. Medical Image Analysis. 2019;54:100–110. doi:10.1016/j.media.2019.02.011.

8. Ben-Cohen A, Mark D, Kovler I, Zur D, Barak A, Iglicki M, et al. Retinal layers segmentation using Fully Convolutional Networkin OCT images. RSIP Vision. 2017.

9. Gorgi Zadeh, S. et al. CNNs Enable Accurate and Fast Segmentation of Drusen in Optical Coherence Tomography International Workshop on Multimodal Learning for Clinical Decision Support. 2017;p.65–73. Available from: https://link.springer.com/chapter/10.1007/978-3-319-67558-9_8.

10. Venhuizen FG, Ginneken Bv, Liefers B, Asten Fv, Schreur V, Fauser S, et al. Deep learning approach for the detection and quantification of intraretinal cystoid fluid in multivendor optical coherence tomography. Biomedical Optics Express. 2018;9(4):1545–1569. doi:10.1364/BOE.9.001545.

11. Chen Z, Li D, Shen H, Mo H, Zeng Z, Wei H. Automated segmentation of fluid regions in optical coherence tomography B-scan images of age-related macular degeneration. Optics & Laser Technology. 2020;122:105830. doi:10.1016/j.optlastec.2019.105830.

12. Roy AG, Conjeti S, Karri SPK, Sheet D, Katouzian A, Wachinger C, et al. ReLayNet: retinal layer and fluid segmentation of macular optical coherence tomography using fully convolutional networks. Biomedical Optics Express. 2017;8(8):3627–3642. doi:10.1364/BOE.8.003627.

13. Alsaih K, Yusoff MZ, Tang TB, Faye I, Mériaudeau F. Deep learning architectures analysis for age-related macular degeneration segmentation on optical coherence tomography scans. Computer Methods and Programs in Biomedicine. 2020;195:105566. doi:10.1016/j.cmpb.2020.105566.

14. Terry L, Trikha S, Bhatia KK, Graham MS, Wood A. Evaluation of Automated Multiclass Fluid Segmentation in Optical Coherence Tomography Images Using the Pegasus Fluid Segmentation Algorithms. Translational Vision Science & Technology. 2021;10(1):27–27. doi:10.1167/tvst.10.1.27.

15. Paszke A, Chaurasia A, Kim S, Culurciello E. ENet: A Deep Neural Network Architecture for Real-Time Semantic Segmentation. 160602147 [cs]. 2016;.

16. Ganjee R, Moghaddam M, Nourinia R An unsupervised hierarchical approach for automatic intra-retinal cyst segmentation in spectral-domain optical coherence tomography images. Medical Physics. 2020;47(10):4872–4884 doi:10.1002/mp.14361.

17. Dabov K, Foi A, Katko V, Egiaza K BM3D Image Denoising with Shape-Adaptive Principal Component Analysis SPARS’09 - Signal Processing with Adaptive Sparse Structured Representations Available from: https://hal.inria.fr/inria-00369582.

18. Cuocolo R, Comelli A, Stefano A, Benfante V et al. Deep Learning Whole-Gland and Zonal Prostate Segmentation on a Public MRI Dataset Journal of Magnetic Resonance Imaging 2021;54(2):452–459 doi:10.1002/jmri.27585

19. Buades A, Coll B, Morel J. Non-Local Means Denoising Image Processing On Line 2011;1:208–212 Available from: https://doi.org/10.5201/ipol.2011.bcm_nlm

20. Rashno A, Koozekanani D, Drayna P, Nazari B, Sadri S, Rabbani H and Parhi K. Fully-Automated Segmentation of Fluid/Cyst Regions in Optical Coherence Tomography Images with Diabetic Macular Edema using Neutrosophic Sets and Graph Algorithms IEEE Transactions on Biomedical Engineering doi:10.1109/TBME.2017.2734058

21. Kermany D. Large Dataset of Labeled Optical Coherence Tomography (OCT) and Chest X-Ray Images. Available from: https://doi.org/10.17632/rscbjbr9sj.3

